# Mfn2 requires Hinge 1 integrity for efficient nucleotide-dependent assembly and membrane fusion

**DOI:** 10.1101/651877

**Authors:** N.B. Samanas, E.A. Engelhart, S. Hoppins

**Author notes:** Corresponding author: Suzanne Hoppins, University of Washington, Department of Biochemistry, 1959 NE Pacific Street, Health Sciences Building, J383, Seattle, WA 98195.

## Abstract

Mitofusins are members of the dynamin-related protein family, large GTPases that harness the energy from nucleotide hydrolysis to remodel membranes. Mitofusins possess four structural domains including two extended helical bundles that are connected by a flexible linker. The role of this linker (Hinge 1) in mitofusin-mediated membrane fusion is not well understood. We have characterized four variants with amino acid substitutions within this region of Mfn2. While a defect was not apparent in cells, a fusion deficiency was observed in vitro, and was rescued by the addition of cytosolic fraction. All four variants had decreased nucleotide-dependent assembly, which was improved by the addition of Bax. Assembly of mitofusins across two membranes was unaffected as formation of the trans complex was similar to wild type for all variants. We further demonstrate that variants with substitutions in both helical bundles are more severely impaired than any single mutant, suggesting that both helical bundles contribute to this function. Our data are consistent with a model where this region contributes to conformational changes that are important for assembly.

## INTRODUCTION

Mitochondrial dynamics have become increasingly recognized as an important indicator of and contributor to both cellular health and death. Mitochondrial shape and their cellular distribution change during the cell cycle, in response to stress, and as part of apoptosis (Tondera et al. 2009; Scorrano 2013; Labbé et al. 2014; Horbay and Bilyy 2016). Mitochondria are trafficked on microtubules and the overall shape and connectivity of the mitochondrial network is maintained or modified through mitochondrial fusion and division, which are mediated by membrane-remodeling large GTPase proteins of the dynamin related protein (DRP) family (Labbé et al. 2014). At steady state, it is estimated that mitochondrial fusion and division events are balanced. When mitochondrial division events exceed fusion events, the network fragments into many small individual mitochondria, and this fragmentation is associated with mitophagy and apoptosis. In contrast, when mitochondrial fusion occurs more frequently than division, the result is a more connected network comprised of longer mitochondria, which is associated with increased ATP production; i.e., during a cellular stress response. The importance of these processes and their regulation is highlighted by the association of dysregulated mitochondrial dynamics with various diseases such as Parkinson’s disease, diabetes, and peripheral neuropathies (Züchner et al. 2004; Vital and Vital 2012; Celardo et al. 2014; Wada and Nakatsuka 2016; Rovira-Llopis et al. 2017).

Mitochondrial DRP-mediated fusion is poorly understood and is mechanistically distinct from both SNARE- and viral-mediated fusion. Fusion DRPs that reside in the outer and inner mitochondrial membranes are mitofusin 1, mitofusin 2 (Mfn1 and Mfn2) and Opa1, respectively. Mfn1 and Mfn2 are functionally related but non-redundant paralogs in mammalian cells (Santel and Fuller 2001; Chen et al. 2003; Eura 2003; Ishihara et al. 2004). Interestingly, mutations in Mfn2, but not Mfn1, are the main cause of the peripheral neuropathy Charcot Marie Tooth Syndrome Type 2A (CMT2A) (Züchner et al. 2004). In CMT2A patients, distal nerve degeneration leads to weakness, sensory loss, gait impairment and foot deformations (Gemignani and Marbini 2001).

Fusion defects associated with some disease-associated variants of Mfn2 can be functionally complemented by Mfn1 (Detmer & Chan 2007b). Consistent with this, a recent report found that expression of exogenous Mfn1 in neurons of a CMT2A mouse model rescued axonal degradation (Zhou et al. 2019). The importance of the interaction between the mitofusin paralogs is further highlighted by the observation that the optimal fusion complex is composed of both Mfn1 and Mfn2 (Hoppins et al. 2011). As with other DRPs, both mitofusins exhibit nucleotide-dependent self-assembly, and the ability to form higher order oligomers has been correlated with fusion activity (Engelhart and Hoppins 2019). Further supporting a role for assembly in DRP-mediated fusion, intermolecular complementation has been observed between non-functional variants of the yeast mitofusin homolog Fzo1 that possess amino acid substitutions in distinct functional domains (Griffin and Chan 2006). Consistently, Fzo1 has been shown to form a large docking ring that generates an extensive area of contact at the interface of two mitochondria (Brandt et al. 2016).

Mitofusins contain four major structural domains including the GTPase domain, two sequential extended helical bundles (HB1 and HB2) connected by flexible loops, and a transmembrane domain (Figure 1). The domain organization is similar to the bacterial DRP (BDLP), which has been structurally characterized (Low and Löwe 2006; Low et al. 2009). In BLDP, the relative positions of these domains changes in different nucleotide states via changes in hinge regions that connect the domains, resulting in either an extended or a closed state (Figure 1A and B). It is hypothesized that mitofusin proteins undergo similar structural rearrangements around hinge regions (Jimah and Hinshaw 2019). Indeed, atomic structures of a minimal GTPase domain construct of Mfn1 demonstrate that relative to the GTPase domain, HB1 can exist in two distinct, nucleotide-dependent conformations due to changes in Hinge 2 (Qi et al. 2016; Cao et al. 2017; Yan et al. 2018). Furthermore, mini-peptides or small molecules thought to alter the stability of the extended or closed state of mitofusin change the overall structure of the mitochondrial network in cells (Franco et al. 2016; Rocha et al. 2018).

**Figure 1.**
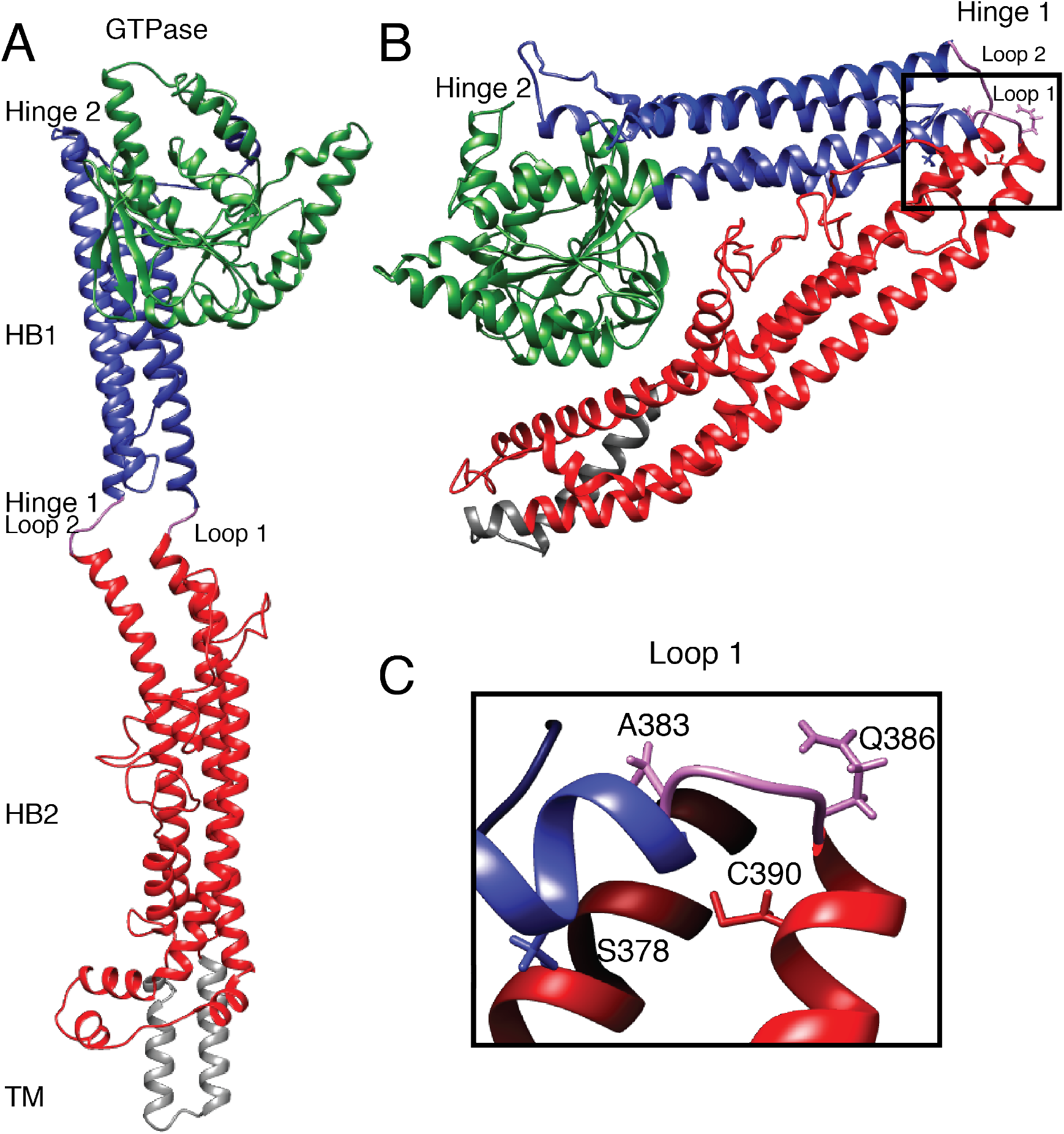
Structural model of the positions of Hinge 1 amino acid substitutions associated with CMT2A. **(A)** Structural model of the predicted extended structure of Mfn2 based on the crystal structure of the structurally related protein BDLP with GMPPNP (PDB 2W6D). The GTPase domain is green, HB1 is blue, HB2 is red, the transmembrane (TM) domain is grey, and Loops 1/ 2 are purple. Structural prediction performed by I-TASSER server (Zhang 2009; Yang and Zhang 2015). **(B)** Structural model of the predicted closed structure of Mfn2 based on the crystal structure of the structurally related protein BDLP with GDP (PDB 2J69). Domains are colored as described in (A). Structural prediction performed by I-TASSER server (Zhang 2009; Yang and Zhang 2015). **(C)** Enlarged view of Loop 1 from Hinge 1 showing the positions of the CMT2A-related amino acids.

The function of the region connecting HB1 and HB2 is relatively unexplored. To identify functionally informative variants of mitofusin and characterize the contribution of this region, we exploited the fact that amino acid substitutions in Mfn2 are associated with human disease. Several of these disease-associated substitutions occur near Hinge 1, consistent with our prediction that this domain is functionally important. To gain insight into the role of Hinge 1 in mitofusin-mediated membrane fusion, we interrogated the mitochondrial fusion activity and biochemical properties of disease-associated hinge variants. Our data indicate that this region is required for optimal mitochondrial fusion activity and efficient nucleotide-dependent assembly of Mfn2.

## RESULTS

### Mfn2 hinge variants restore reticular mitochondrial morphology in Mfn2-null cells

We set out to characterize four mutant variants of Mfn2 with amino acid substitutions within or adjacent to Loop 1 of Hinge 1: S378P, A383V, Q386P, and C390F (Figure 1C and Table S1). We began by analyzing mitochondrial morphology in stable cell lines expressing mutant versions of Mfn2. To create these lines, we introduced either wild type or mutant *MFN2* with a C-terminal 3xFLAG tag into Mfn2-null mouse embryonic fibroblasts (MEFs) using retroviral transduction (Chen et al. 2003). Clonal populations expressing Mfn2 at near-endogenous levels were selected for characterization (Figure S1).

In wild type MEFs, mitochondria were in reticular networks where most of the mitochondria were longer than 2.5 µm (Figure 2, Mfn1^+/+^Mfn2^+/+^). Mfn2-null cells transduced with an empty vector, conversely, contained mainly fragmented individual mitochondria less than 2.5 µm in length (Figure 2, vector), which is consistent with published observations of Mfn2-null cells (Chen et al. 2003). Expression of wild type Mfn2 in Mfn2-null cells restored fusion activity and a reticular mitochondrial network in about 80% of cells (Figure 2, Mfn2^WT^). Somewhat surprisingly, all Mfn2 mutant variants examined here (Mfn2^S378P^, Mfn2^A383V^, Mfn2^Q386P^, and Mfn2^C390F^) also restored a reticular mitochondrial network in Mfn2-null cells to a similar extent as Mfn2^WT^. These results indicate that all four of these Mfn2 variants are stable and well-folded and support robust fusion activity in Mfn2-null fibroblasts. This is consistent with previously reported data that showed subsets of CMT2A associated mutations restored fusion activity (Detmer & Chan 2007; Engelhart & Hoppins 2019).

**Figure 2.**
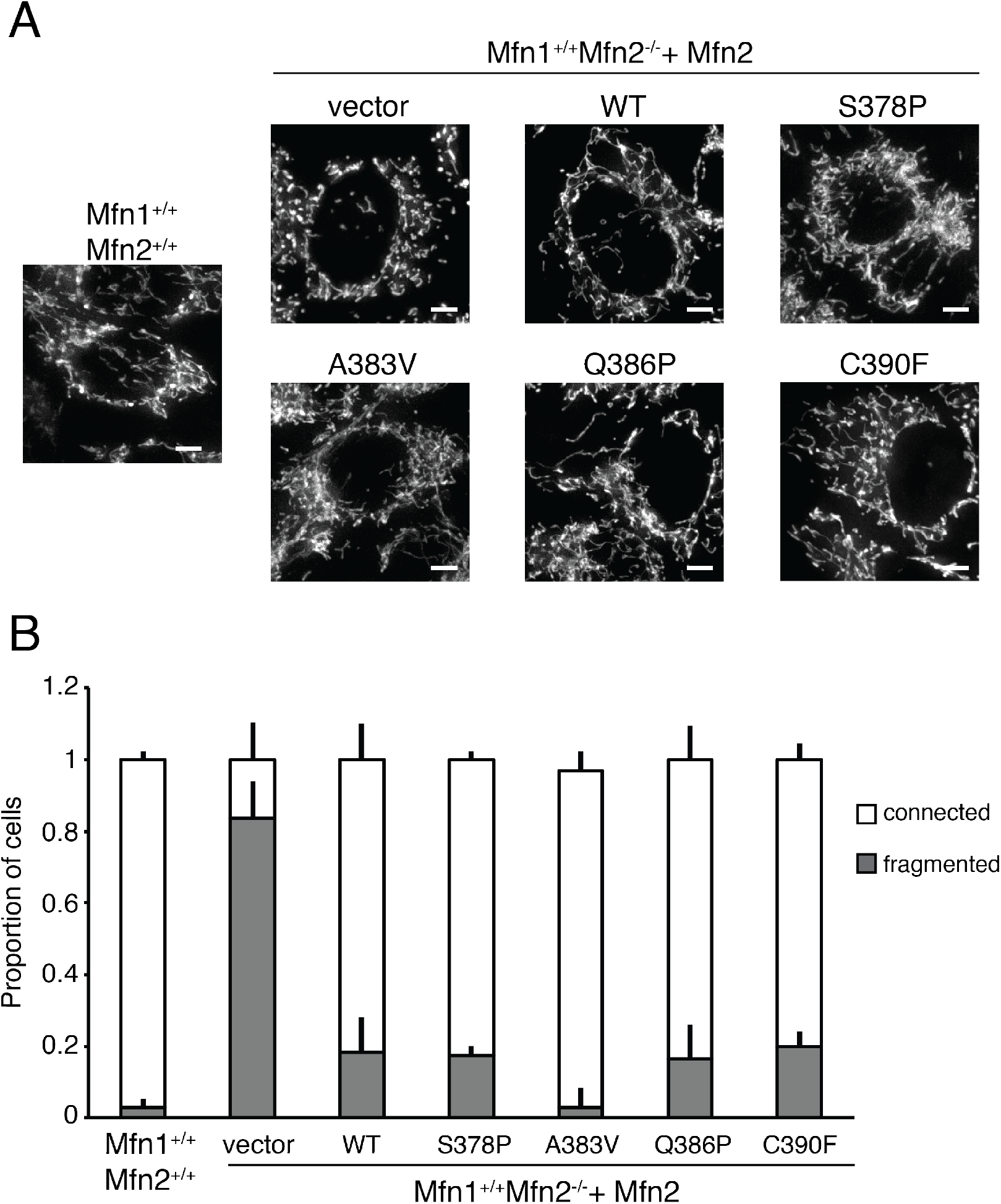
Mfn2 hinge variants support mitochondrial fusion when expressed in Mfn2-null cells. **(A)** Representative images of mitochondrial networks in wild type (Mfn1^+/+^Mfn2^+/+^) or Mfn2-null (Mfn1^+/+^Mfn2^-/-^) mouse embryonic fibroblasts expressing the indicated Mfn2 variant. Mitochondria were stained with Mitotracker Red CMXRos and visualized by fluorescence microscopy. Images represent a maximum intensity projection. Scale bars = 5 μm. **(B)** Quantification of mitochondrial morphology in cells represented in **(A)** Error bars indicate mean + standard deviation from three blinded experiments (n ≧ 100 cells per population per experiment).

### Mfn2 hinge variants have an in vitro mitochondrial fusion defect

We reasoned that these substitutions may result in biochemical changes to the protein that were masked in the context of the cell, where many factors modulate the structure of the mitochondrial network. Therefore, we proceeded to characterize these mutant forms of Mfn2 in the context of isolated mitochondria. To quantify the mitochondrial fusion activity of the Mfn2 mutant variants in the absence of cytosolic factors, we utilized a cell-free mitochondrial fusion assay. Mitochondria isolated from cells expressing RFP or CFP targeted to the mitochondrial matrix were mixed, incubated in fusion buffer and then imaged by fluorescence microscopy. Fusion events were scored as the overlap of the two fluorophores in three dimensions. All assays were performed in parallel with mitochondria isolated from wild type cells, and data is expressed as a proportion of wild type controls. Mitochondria isolated from Mfn2-null cells transduced with empty vector fused at a much lower frequency than wild type controls (Figure 3A). Fusion of mitochondria isolated from the clonal population of Mfn2-null cells expressing Mfn2^WT^ was similar to wild type controls (Figure 3A), consistent with the restoration of the mitochondrial morphology in cells. To quantify the fusion activity of the Mfn2 hinge mutants in vitro, mitochondria were isolated from each of the clonal populations described above. In each case, the mitochondrial fusion activity was significantly lower than wild type controls (Figure 3A).

**Figure 3.**
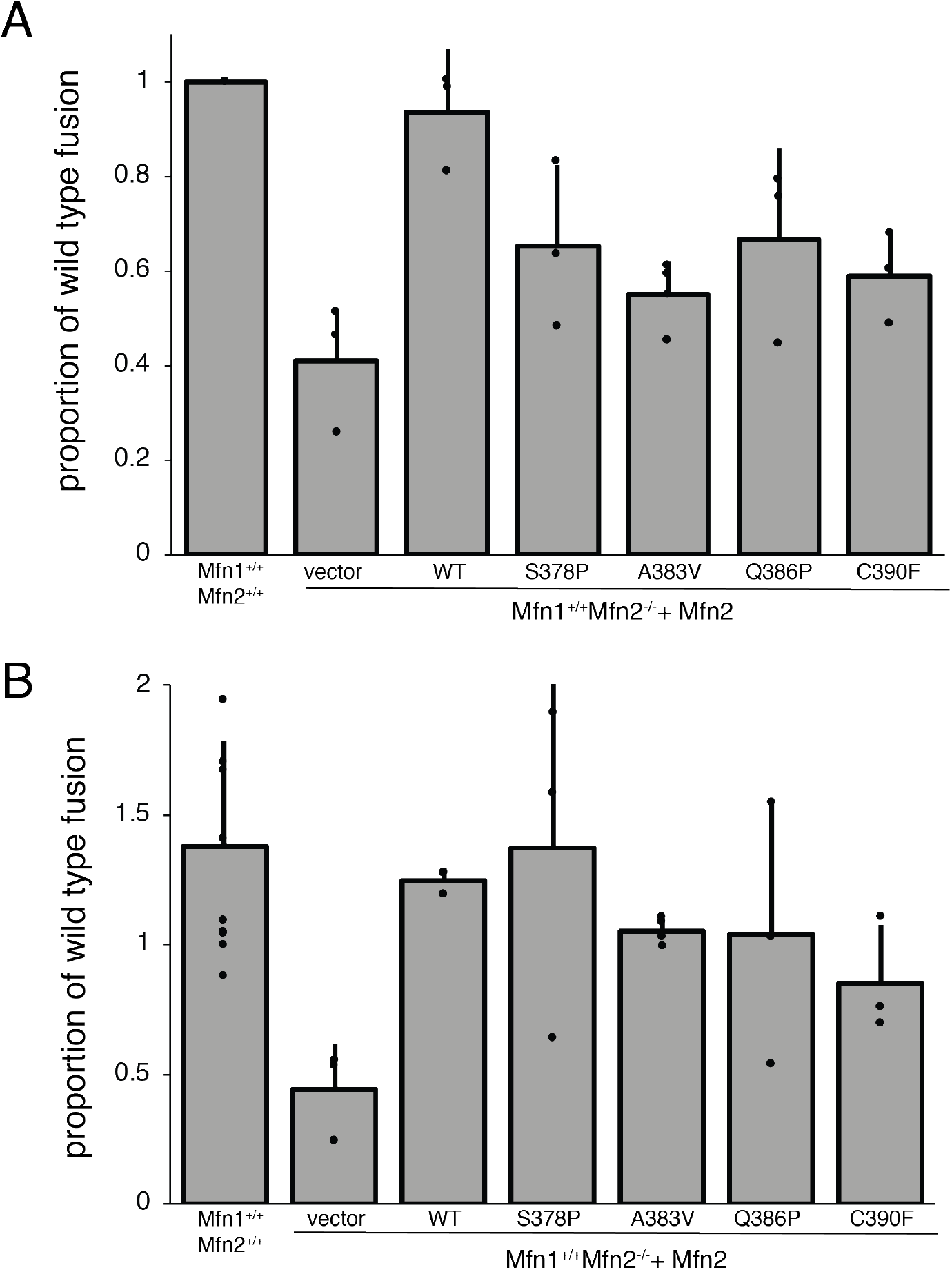
Mitochondrial in vitro fusion assay reveals a defect for all Mfn2 hinge variants. **(A)** Mitochondria were isolated from wild type cells or clonal populations of Mfn2-null cells either transduced with empty vector or expressing the indicated Mfn2 variant were subject to in vitro fusion conditions at 37°C for 60 minutes. The data are represented as relative to wild type controls performed in parallel. Error bars indicate mean + standard deviation from at least three independent experiments and the statistical significance were determined by paired t-test analysis (*P<0.05). **(B)** Mitochondrial in vitro fusion assay performed as in (A) except with the addition of cytosol-enriched fraction to the reaction buffer. Data are represented as relative to wild type controls performed in parallel without cytosol added. Error bars indicate mean + standard deviation from at least three independent experiments

The in vitro mitochondrial fusion data revealed that amino acid substitutions in this hinge region diminish the fusion activity of Mfn2. Given that we could not detect a mitochondrial fusion defect in cells, we predicted that a cytosolic factor could be enhancing fusion activity of the mutant variants. To test this in vitro, we performed the mitochondrial fusion assay in the presence of cytosol from wild type MEFs. As has been previously reported, the addition of the cytosol-enriched fraction to wild type mitochondria moderately stimulated fusion activity (Hoppins et al. 2011). In contrast, the cytosol-enriched fraction did not alter the fusion activity of mitochondria isolated from Mfn2-null cells, indicating that Mfn2, but not Mfn1, is regulated by the cytosolic factor (Figure 3B). The addition of cytosol also increased the fusion activity of mitochondria that possess the Mfn2 hinge variants similarly to wild type controls (Figure 3B). These data indicate that the fusion defect associated with the hinge mutant variants can be compensated for by cytosolic factors in cells.

### Mfn2 hinge mutant variants interact with Mfn1 in cis and in trans

Mfn1 and Mfn2 physically interact in the same membrane, in cis, and across two membranes, in trans, as measured by co-immunoprecipitation (Chen et al. 2003; Scott A Detmer and Chan 2007b; Engelhart and Hoppins 2019). To determine if the Mfn2 hinge mutant variants interact with Mfn1 in cis or in trans, we tested whether Mfn1 would co-immunoprecipitate with Mfn2-FLAG. To distinguish between cis and trans, we mixed mitochondria that possess endogenous Mfn1 and Mfn2-FLAG with mitochondria that possess Mfn1-EGFP and endogenous Mfn2 (Figure 4A). In this way, three unique interactions with Mfn2-FLAG (Figure 4, filled arrowhead) could be assessed: (1) endogenous Mfn1 in cis (Figure 4, black arrow), (2) Mfn1-EGFP in trans (Figure 4, open arrowhead), and (3) endogenous Mfn2 in trans (Figure 4, white arrow). These reactions were performed in the presence of the GTP transition state mimic GDP-BeF_3_, which has been shown to stabilize an interaction between mitofusin molecules and promote a tethering interaction, or BeF_3_ alone as a negative control (Qi et al. 2016; Yan et al. 2018; Engelhart and Hoppins 2019). As expected, wild type Mfn2-FLAG immunoprecipitated both endogenous Mfn1 and Mfn1-EGFP (Figure 4). The amount of endogenous Mfn1 interacting in cis (Figure 4, black arrow) is not significantly different in the presence or absence of the transition state mimic. In contrast, the trans interaction is more robust in the presence of GDP-BeF_3_ compared to BeF_3_ alone (Figure 4, open arrowhead). Together, these data indicate that only the trans interaction is highly dependent on the nucleotide binding state of the mitofusin proteins. Each of the four Mfn2 hinge variants also immunoprecipitated Mfn1 in cis and in trans (Figure 4), indicating that there is no defect in the physical interaction with Mfn1 in either context. Interestingly, even in the presence of GDP-BeF_3_, we could not detect an interaction in trans between Mfn2-FLAG and endogenous Mfn2 (Figure 4, white arrow), which suggests that Mfn2 does not form a robust homotypic trans complex.

**Figure 4.**
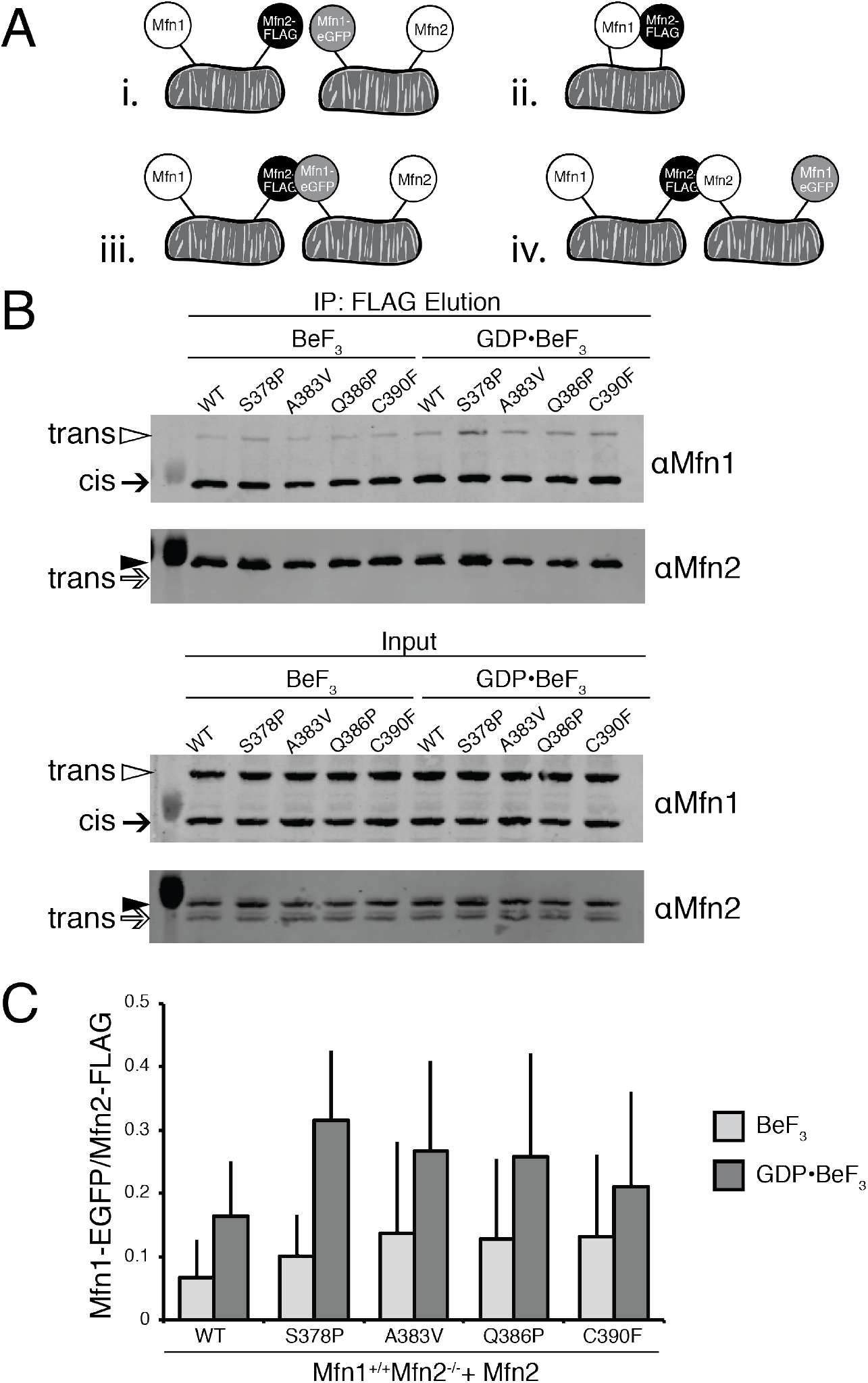
Mfn2 hinge variants interact with Mfn1 in cis and trans. **(A)** Schematic of the differential epitope labeling utilized in the co-immunoprecipitation assay. **(B)** Mitochondria were isolated from a clonal population of Mfn1-null cells expressing Mfn1^WT^-EGFP at endogenous levels (Mfn1^WT^Mfn2^+/+^), and clonal populations of Mfn2-null cells expressing the indicated Mfn2-FLAG variants. Mitochondria that possess Mfn1-EGFP and Mfn2 were combined with mitochondria that possess Mfn1 and Mfn2-FLAG and these mixtures were incubated with BeF_3_ in the absence or presence of GDP. Following lysis, immunoprecipitation was performed with α-FLAG magnetic beads. Proteins eluted from the beads were subjected to SDS-PAGE and immunoblotting with α-Mfn1 and α-Mfn2, as indicated. Arrows indicate endogenous Mfn1 (black) and Mfn2 (white); arrowheads indicate Mfn1-EGFP (white) and Mfn2-FLAG (black), respectively. Input represents 3% of the input and elution represents 37.5% of the immunoprecipitated protein. **(C)** Quantification of the percentage of Mfn1-EGFP in the elution compared to Mfn2-FLAG is shown as the mean + standard deviation of three independent experiments.

### Nucleotide-dependent self-assembly is diminished in Mfn2 hinge mutant variants

Defects in mitofusin assembly correlate with reduced rates of mitochondrial fusion in vitro (Engelhart and Hoppins 2019). To determine the capacity of the Mfn2 hinge variants to assemble, we utilized blue native gel electrophoresis (BN-PAGE). Mitochondria were left untreated or were incubated with either GTP or the non-hydrolyzable analog GMPPNP. Mitochondria were then lysed and separated by BN-PAGE and subject to Western blot analysis. When mitochondria were untreated, Mfn2^WT^ migrated mostly in assemblies that approximately correspond in size to a dimer (Figure 5A and C, arrow), with some protein migrating as larger assemblies (Figure 5A and C, arrowheads). When these mitochondria were instead incubated with nucleotide, the ratio of dimer to larger assemblies decreased, consistent with nucleotide-dependent assembly of Mfn2 into higher-order oligomers. Specifically, following incubation with GTP, more Mfn2^WT^ migrated in two larger assemblies, with a notable increase in the ∼450 kDa oligomer (Figure 5A, open arrowhead). Incubation of mitochondria with GMPPNP also promoted higher-order assembly with enhanced stability of a ∼320 kDa oligomer and some protein migrating as the ∼450 kDa oligomer (Figure 5A, closed and open arrowheads, respectively). Together, these data indicate that Mfn2 exists primarily as a dimer in the mitochondrial outer membrane and assembles into at least two larger oligomeric species in a nucleotide-dependent manner.

**Figure 5.**
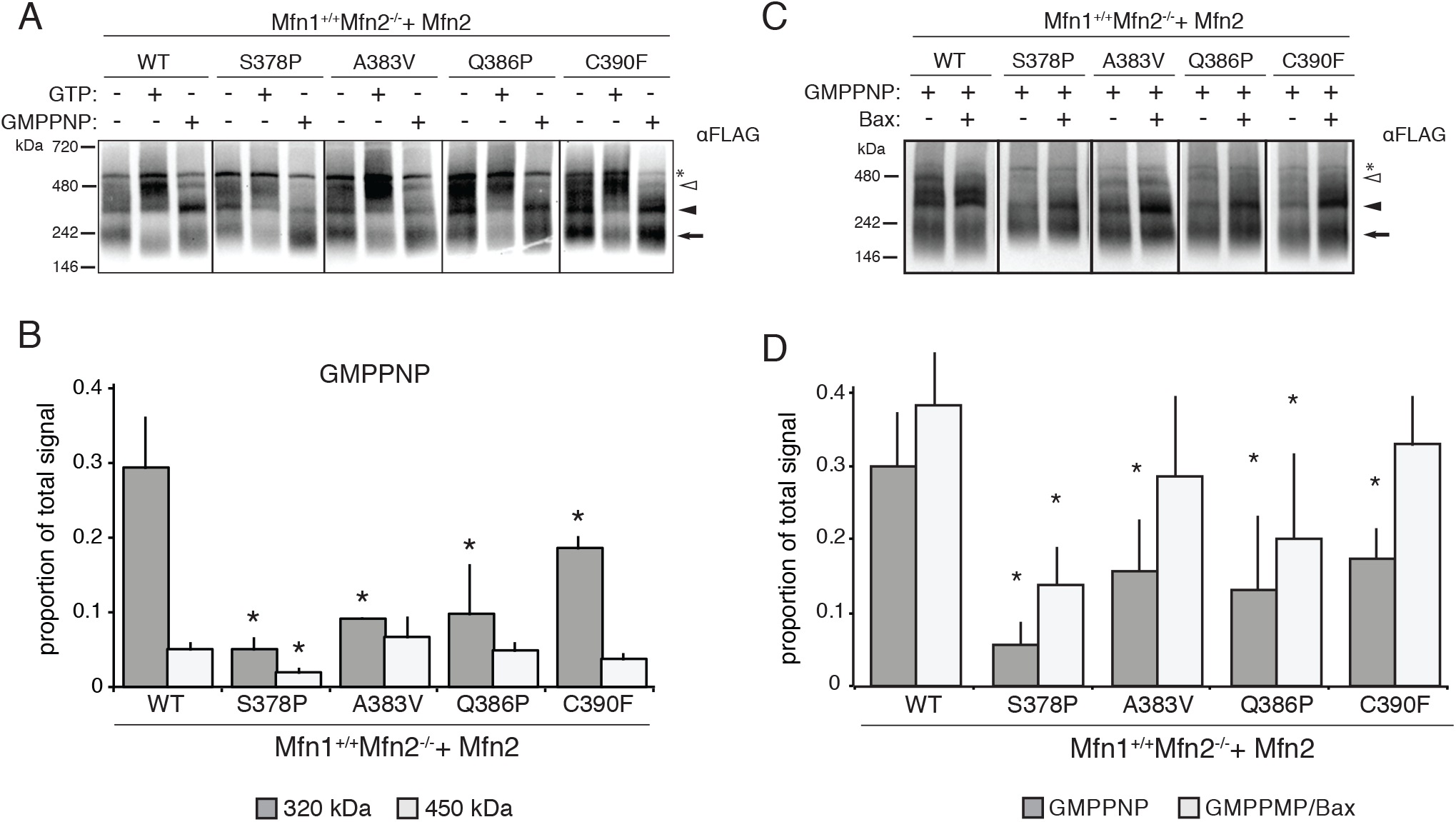
Mfn2 hinge variants have altered nucleotide-dependent assembly. **(A)** Mitochondria were isolated from clonal populations of Mfn2-null cells expressing the indicated Mfn2 variant. Mitochondria were either untreated or incubated with 2mM GTP or 2mM GMPPNP as indicated before lysis and separation by BN-PAGE followed by immunoblotting with anti-FLAG antibody. Arrow represents predicted dimer, closed arrowhead indicates ∼320 kDa species, and open arrowhead indicates ∼450 kDa species. Asterisk (*) indicates nonspecific signal. **(B)** Quantification of proportion of the total protein observed in the 320kDa or 450kDa band after treatment with GMPPNP (filled arrowhead). Error bars indicate mean + standard deviation from at least three independent experiments and the statistical significance were determined by paired t-test analysis between the indicated data and wild type (*P<0.05). **(C)** Mitochondria were prepared as in (A) and incubated with 2mM GMPPNP with or without 1μM recombinant purified Bax. Arrow represents predicted dimer, closed arrowhead indicates ∼320 kDa species, and open arrowhead indicates ∼450 kDa species. Asterisk (*) indicates nonspecific signal. **(D)** Quantification of proportion of the total protein observed in the 320kDa band. Error bars indicate mean + standard deviation from at least three independent experiments and the statistical significance were determined by paired t-test analysis between the indicated data and wild type (*P<0.05).

In untreated mitochondria, all of the Mfn2 hinge variants migrated similarly to wild type, with most of the protein in the dimer (Figure 5A). In the presence of GTP, the mutants also assembled similar to wild type, with the exception of Mfn2^S378P^, which had less protein migrating as the 450 kDa oligomer compared to wild type. Each of the hinge mutants displayed altered assembly relative to wild type Mfn2 in the presence of GMPPNP (Figure 5B). Compared to Mfn2^WT^, Mfn2^S378P^, Mfn2^A383V^, Mfn2^Q386P^, and Mfn2^C390F^ all showed a significantly decreased amount of protein migrating as the 320 kDa oligomer (Figure 5B). These data indicate that amino acid substitutions in this hinge region prevent the stable assembly of Mfn2 into this oligomeric species in the presence of the non-hydrolyzable GTP analog. Given that these mutant variants also exhibit an in vitro mitochondrial fusion defect, we hypothesize that nucleotide dependent assembly contributes to efficient Mfn2-mediated membrane fusion. Our data would further predict that cytosolic factors, which improve in vitro fusion activity, could facilitate Mfn2 assembly. To test this, we assessed Mfn2 nucleotide-dependent assembly in the presence of recombinant purified Bax, which has been previously demonstrated to promote Mfn2-dependent mitochondrial fusion and Mfn2 assembly (Karbowski et al. 2006; Hoppins et al. 2011). In the presence of Bax, we observe more Mfn2^WT^ in the 320 kDa oligomer, consistent with Bax playing a role in Mfn2 assembly (Figure 5C and 5D). Furthermore, Mfn2^S378P^, Mfn2^A383V^, Mfn2^Q386P^, and Mfn2^C390F^ also had increased abundance of the 320 kDa oligomer in the presence of Bax (Figure 5C and 5D). Together, these data support the conclusion that impaired nucleotide-dependent assembly results in diminished in vitro fusion activity and that cytosolic factors such as Bax compensate for these defects in cells.

### Double hinge mutations reveal that HB1 and HB2 work together in mitochondrial fusion

Our data indicate that this region of Mfn2 is important for Mfn2 fusion activity. Based on structural models, S378 is predicted to be located in HB1 while C390 is part of HB2. We considered the possibility that these two distinct structural domains function in cooperatively to control the conformational state of Mfn2. If this is the case, a variant of Mfn2 with amino acid substitutions in both HB1 and HB2 would be predicted to have a more severe functional defect than variants with two substitutions in the same structural domain. To test this hypothesis, we used Mfn2^L710P^, a disease-associated variant in HB2 that is located near Loop 2 in Hinge 1 (Verhoeven et al. 2006) (Figure 1, Figure S2). This amino acid substitution has been characterized in Mfn1, Mfn1^L691P^, which had some fusion activity in Mfn1-null cells (Koshiba et al. 2004). We generated double mutant variants of Mfn2 with this mutation and either Mfn2^S378P^, which is located in HB1 or Mfn2^C390F^, which is located in HB2 (Mfn2^S378P/L710P^ and Mfn2^C390F/L710P^, respectively).

To determine the fusion activity of these double mutant variants, we utilized retroviral transduction to express Mfn2-mNeonGreen (Mfn2-NG) in Mfn2-null MEFs which has previously been shown to restore fusion activity (Engelhart and Hoppins 2019). The mitochondrial morphology was scored in cells expressing Mfn2, as assessed by colocalization of Mfn2-NG with MitoTracker Red. As expected, Mfn2^WT^, Mfn2^S378P^ and Mfn2^C390F^ restored the reticular mitochondrial network, which recapitulated our observations from the clonal populations described above (Figure 6, compare to Figure 2). Expression of Mfn2^L710P^ also restored a reticular network in 65% of cells (Figure 6), indicating a mild defect in mitochondrial fusion, consistent with previous reports (Koshiba et al. 2004).

**Figure 6.**
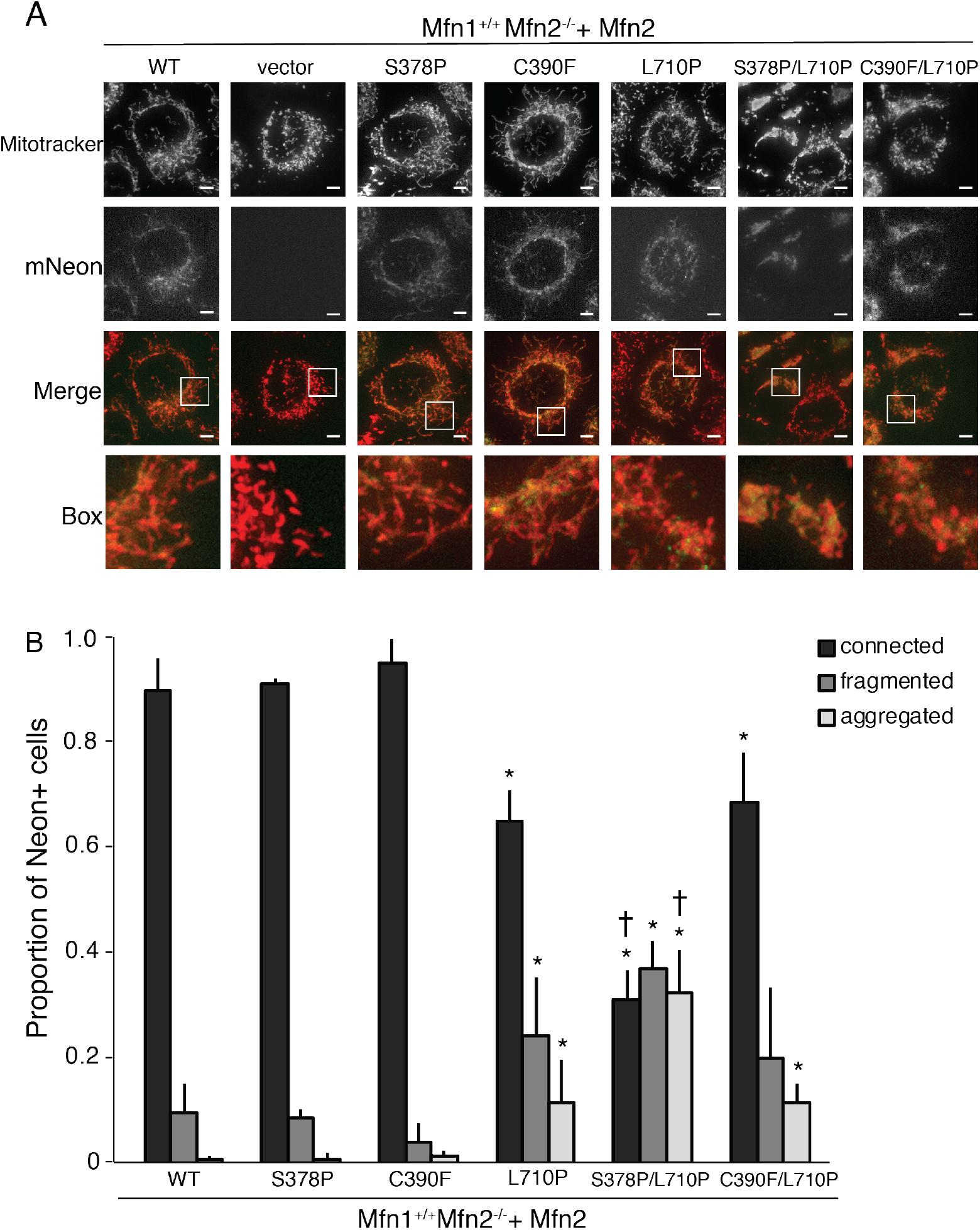
Mfn2 variants with substitutions in both HB1 and HB2 are defective for fusion in Mfn2-null cells. **(A)** Representative images of mitochondrial networks in Mfn2-null (Mfn1^+/+^Mfn2^-/-^) mouse embryonic fibroblasts expressing the indicated Mfn2-mNeonGreen variant. Mitochondria were stained with Mitotracker Red CMXRos and visualized by fluorescence microscopy. Images represent a maximum intensity projection. Scale bars = 5 μm. **(B)** Quantification of mitochondrial morphology of cells represented in **(A).** Error bars indicate mean + standard deviation from at least three independent experiments and the statistical significance were determined by paired t-test analysis between the indicated data and wild type (*P<0.05) or between the indicated data and Mfn2^S378P/L710P^ (†P<0.05).

In Mfn2-null cells expressing the HB2 double mutant Mfn2 ^C390F/L710P^, the mitochondrial morphology was comparable to that in cells expressing Mfn2^L710P^ alone (Figure 6). In contrast, when both HB1 and HB2 possess an amino acid substitution, there is significantly less mitochondrial fusion. Specifically, in Mfn2-null cells expressing Mfn2 ^S378P/L710P^, most cells possessed a mitochondrial network that was either fragmented or in fragmented aggregates, and only 30% of cells had a reticular mitochondrial network. Together, these results support the conclusion that HB1 and HB2 provide unique and separate contributions to support Mfn2-mediated membrane fusion.

## DISCUSSION

In this study, we performed an in-depth characterization of disease-associated amino acid substitutions located within Hinge 1 of Mfn2, which connects HB1 and HB2. In other DRPs, hinges mediate conformational changes that are required for membrane remodeling activity. The results presented here are consistent with structural models that suggest this region functions as a hinge and for the first time connects Hinge 1 integrity to nucleotide-dependent assembly of the mitofusins. These molecular defects also correspond to a decrease in mitochondrial fusion efficiency in vitro. Both of these defects were improved by the addition of cytosolic factors, which indicates that mitofusin function is subject to complex regulation in cells.

Our data indicate that Mfn2 exists primarily as a dimer in the mitochondrial outer membrane and this form is not altered by the amino acid substitutions tested here. Furthermore, the Mfn2 variants are able to interact with Mfn1 on the same and adjacent mitochondria in a nucleotide dependent manner similarly to wild type Mfn2. Together, these data suggest that these amino acid substitutions do not significantly alter the structure of Mfn2. We demonstrate that GTP stabilizes a 450 kDa oligomer while the non-hydrolyzable analog stabilizes a 320 kDa oligomer. In the presence of GTP, most of the migration pattern of the hinge variants is similar to wild type and each can form both dimers and 450 kDa oligomers. This indicates that these variants are not unable to fold or oligomerize. In contrast, the hinge mutants do not readily form the 320 kDa state adopted when Mfn2 is bound to GMPPNP. Given that all Mfn2 hinge mutants co-immunoprecipitate Mfn1 as efficiently as wild type, interaction with Mfn1 is not likely to play a role in this assembly. Indeed, previously published data from our lab suggest that the BN-PAGE assemblies are homo-oligomers (Engelhart and Hoppins 2019). These data are consistent with the hypothesis that the nucleotide binding state of Mfn2 alters the conformational state of the protein, which results in the preferential formation of distinct oligomeric species. Therefore, the reduced assembly observed for the hinge mutants could be due to aberrant structural rearrangements of HB1 and HB2 as mediated by Hinge 1. Our data further implicate Bax in the formation or stability of these oligomers.

Functional complementation has been observed between two non-functional alleles of Fzo1, the mitofusin homolog (Griffin and Chan 2006). These data indicate that different molecules in the fusion complex can contribute different properties. The amino acid substitutions tested here are predicted to be within either HB1 (S378), HB2 (C390) or a flexible loop between the two (A383V, Q386P). Given that each individual amino acid substitution was associated with a similar partial loss of function, we considered that these structural domains may work cooperatively to evoke nucleotide-dependent conformational changes. To test this, we assessed the fusion activity of variants with two amino acid substitutions, either in the same domain (HB2, C390F & L710P) or in different domains (HB1, S378P & HB2, L710P). When amino acid substitutions were both in HB2, the fusion activity in cells was more substantial than when the substitutions were in HB1 and HB2. Therefore, our data indicate that distinct structural domains provide unique contributions to Mfn2 fusion activity within the same molecule. As a whole, the data presented here are consistent with the hypothesis that different nucleotide-dependent conformations support either the formation or stabilization of different assembly states and that the integrity of the hinge that connects HB1 and HB2 plays a role in this process. We have further shown that this assembly is supported not only by intramolecular interactions, but also by regulating cytosolic factors such as Bax.

## SUPPLEMENTAL FIGURES

**Table S1.** Clinical characterization of the Hinge 1 Loop 1 CMT2A mutations in this study.

| **Mutation** | **Age at onset** | **Severity of symptoms** | **Other** | **Reference** |
| --- | --- | --- | --- | --- |
| S378P | 7 | Mild compared to others in the same study |  | Brockmann, et al. 2007 |
| A383V | various | Various; apparent incomplete penetrance | 2 unrelated families with same mutation | Muglia, et al., 2007 |
| Q386P | 1.5 | Severe | De novo mutation | Verhoeven, et al., 2006 |
| C390F | 1 | Severe | De novo mutation | Feely et al., 2011 |

**Figure S1.**
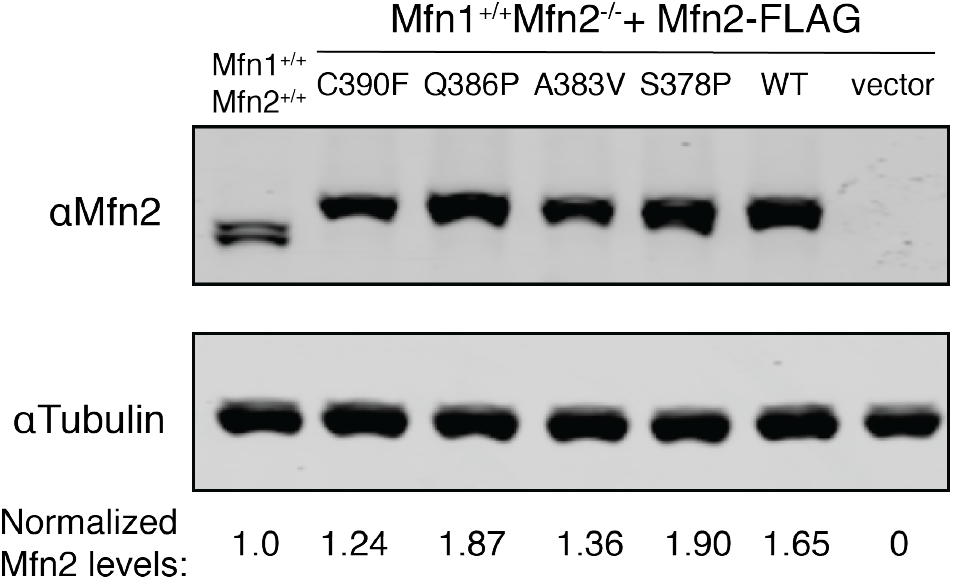
Mfn2 protein expression in MEF clonal populations. Whole-cell lysates prepared from the indicated cell lines were subject to SDS-PAGE and immunoblotting with *α*-Mfn2 and *α* -tubulin.

**Figure S2.**
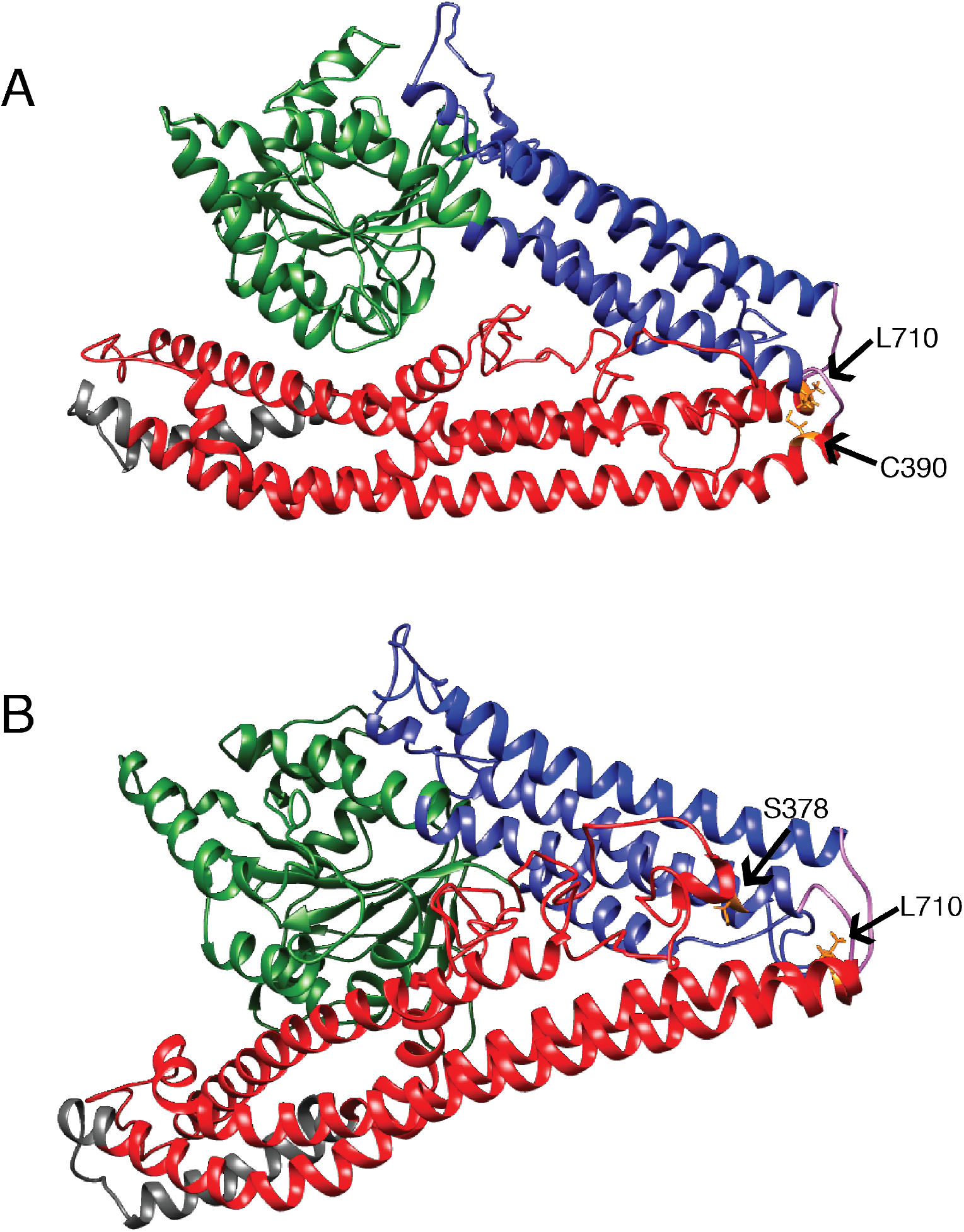
Structural model of the positions of hinge amino acid substitutions associated with CMT2A. Structural models of the predicted closed structure of Mfn2 based on the crystal structure of the structurally related protein BDLP with GDP (PDB 2J69). The GTPase domain is green, HB1 is blue, HB2 is red, the transmembrane (TM) domain is grey, and Loops 1/ 2 are purple. **(A)** The positions of C390 and L710 are indicated in orange and labeled. **(B)** The positions of S378 and L710 are indicated in orange and labeled. Structural prediction performed by I-TASSER server (Zhang 2009; Yang and Zhang 2015).

## MATERIALS AND METHODS

### Cell culture

All cells were grown at 37°C and 5% CO_2_ and cultured in DMEM (Thermo Fisher Scientific) containing 1X GlutaMAX (Thermo Fisher Scientific) with 10% FBS (Seradigm) and 1% penicillin/streptomycin (Thermo Fisher Scientific). Mouse embryonic fibroblasts cells (Mfn wildtype and Mfn2-null) were purchased from ATCC.

### Retroviral transduction and generation of clonal populations

Plat-E cells (Cell Biolabs) were maintained in complete media supplemented with 1 μg/mL puromycin and 10 μg/mL blasticidin and plated at approximately 80% confluency the day prior to transfection. Plat-E cells were transfected with FuGENE™ HD (Promega) and transfection regent was incubated overnight before a media change. Viral supernatants were collected at approximately 48, 56, 72, and 80 hours post transfection and incubated with MEFs in the presence of 8 mg/ml polybrene. Approximately 16 hours after the last viral transduction, MEF cells were split and selection was added if needed (1 μg/mL puromycin or 200 μg/mL hygromycin).

Clonal populations were generated by plating cells at very low density and clones were collected onto sterile filter paper dots soaked in trypsin. Following expansion, whole cell extract from clonal populations were screened by western blot analysis for mitofusin against wildtype controls.

### Transfection and microscopy

All cells were plated in No. 1.5 glass-bottomed dishes (MatTek). Mouse embryonic fibroblasts were incubated with 0.1 μg/mL Mitotracker Red CMX Ros (Invitrogen) for 15 minutes at 37°C with 5% CO_2_, washed and incubated with complete media for at least 45 minutes prior to imaging. MEFs were imaged at 37°C with 5% CO_2_. A Z-series with a step size of 0.3 μm was collected with a Nikon Ti-E widefield microscope with a 63X NA 1.4 oil objective (Nikon), a solid-state light source (Spectra X, Lumencor), and an sCMOS camera (Zyla 5.5 Megapixel). Each cell line was imaged by a blinded researcher on at least three separate occasions (n > 100 cells per experiment).

### Image analysis

Images were deconvolved using 8-15 iterations of 3D Landweber deconvolution. Deconvolved images were then analyzed using Nikon Elements software. Maximum intensity projections were created using Photoshop (Adobe). Mitochondrial morphology was scored as follows: reticular indicates that fewer than 30% of the mitochondria in the cell were fragments (fragments defined as mitochondria less than 2 μm in length); fragmented indicates that most of the mitochondria in the cell were less than 2 μm in length; aggregated indicates fragmented mitochondria that were not distributed throughout the cytosol.

### Preparation of mitochondria or cytosol-enriched fraction

For each experiment, three to five 15 cm plates each of MEFs were grown to ∼90% confluency. Cells were harvested by cell scrapping, pelleted, and washed in mitochondrial isolation buffer (MIB) (0.2 M sucrose, 10 mM Tris-MOPS [pH 7.4], 1 mM EGTA). The cell pellet was resuspended in one cell pellet volume of cold MIB, and cells were homogenized by 10 to 14 strokes on ice with a Kontes Potter-Elvehjem tissue grinder set at 400 RPM. The homogenate was centrifuged (500 × *g*, 5 min, 4°C) to remove nuclei and unbroken cells, and homogenization of the pellet fraction was repeated followed by centrifugation at 500 × *g*, 5 min, 4°C. The supernatant fractions were combined and centrifuged again at 500 × *g*, 5 min, 4°C to remove remaining debris. The supernatant was transferred to a clean microfuge tube and centrifuged (7400 × *g*, 10 min, 4°C) to pellet a crude mitochondrial fraction. The post-mitochondrial supernatant fraction was saved as the cytosol-enriched fraction. The crude mitochondrial pellet was resuspended in a small volume of MIB. Protein concentration of fractions was determined by Bradford assay (Bio-Rad Laboratories).

### In vitro mitochondrial fusion

An equivalent mass (12.5 μg) of mtTagRFP and mtCFP mitochondria were mixed, washed in 500uL MIB and concentrated by centrifugation (7400 × *g*, 10 min, 4°C). Following a 10 min incubation on ice, the supernatant was removed and the mitochondrial pellet was resuspended in 10 μl fusion buffer (20 mM PIPES-KOH [pH 6.8], 150 mM KOAc, 5 mM Mg(OAc)_2_, 0.4 M sorbitol, 0.12 mg/ml creatine phosphokinase, 40 mM creatine phosphate, 1.5 mM ATP, 1.5 mM GTP) or 10 μl cytosol-enriched buffer (2.5 μL of the cytosol-enriched fraction obtained from WT MEFs and 7.5 μL fusion buffer). Fusion reactions were incubated at 37°C for 60 minutes.

### Analysis of mitochondrial fusion

Mitochondria were imaged on depression microscope slides by pipetting 4μL fusion reaction onto a 3% low-melt agarose bed, made in modified fusion buffer (20 mM PIPES-KOH [pH 6.8], 150 mM KOAc, 5 mM Mg(OAc)_2_, 0.4 M sorbitol). A Z-series of 6 0.2 μm steps was collected with a Nikon Ti-E widefield microscope with a 100X NA 1.4 oil objective (Nikon), a solid state light source (Spectra X, Lumencor), and a sCMOS camera (Zyla 5.5 Megapixel). For each condition tested, mitochondrial fusion was assessed by counting ≥ 300 total mitochondria per condition from ≥ 4 images per condition (50 – 200 mitochondria per image collected), and fusion was scored by colocalization of the red and cyan fluorophores in three dimensions.

### BN-PAGE

Isolated mitochondria (15 – 30 μg) were incubated with or without 2 mM nucleotide (GTP or GMPPNP) and 1μM purified recombinant Bax as indicated in modified mitochondrial isolation buffer (0.2 M sucrose, 10 mM Tris-MOPS [pH 7.4], 1 mM EGTA, 5 mM Mg(OAc)_2_, 50mM KOAc, 1X HALT protease inhibitor (Thermo Scientific), 0.5mM phenylmethylsulfonyl fluoride (PMSF)) at 37°C for 30 minutes. Mitochondria were then lysed in 1% w/v digitonin, 50 mM Bis-Tris, 50 mM NaCl, 10% w/v glycerol, 0.001% Ponceau S; pH 7.2 for 15 minutes on ice. Lysates were centrifuged at 16,000 x *g* at 4° C for 30 minutes. The cleared lysate was mixed with Invitrogen NativePAGE^TM^ 5% G-250 Sample Additive to a final concentration of 0.25%. Samples were separated on a Novex™ NativePAGE™ 4 - 16% Bis-Tris Protein Gels (Invitrogen) <u>at 4</u>°<u>C.</u> Gels were run at 40 V for 30 minutes then 100 V for 30 minutes with dark cathode buffer (1X NativePAGE^TM^ Running Buffer (Invitrogen), 0.02% (w/v) Coomassie G-250). Dark cathode buffer was replaced with light cathode buffer (1X NativePAGE^TM^ Running Buffer (Invitrogen), 0.002% (w/v) Coomassie G-250) and the gel was run at 100 V for 30 minutes and subsequently at 250 V for 60-75 minutes until the dye front ran off the gel. After electrophoresis was complete, gels were transferred to PVDF membrane (Bio-Rad Laboratories) at 30 volts for 16 hours in transfer buffer (25 mM Tris, 192 mM glycine, 20% methanol). Membranes were incubated with 8% acetic acid for 15 minutes and washed with H_2_O for 5 minutes. Membranes were dried at 37°C for 20 minutes and then rehydrated in 100% methanol and washed in H_2_O. Membranes were blocked in 4% milk for 20 minutes and were probed with anti-FLAG (Sigma) for 4 hours at room temperature or overnight at 4°C. Membranes were incubated with HRP-linked secondary antibody (Cell Signaling Technology) at room temperature for 1 hour. Membranes were developed in SuperSignal Femto ECL reagent (Thermo Fisher Scientific) for 5 minutes and imaged on iBright Imaging System (Thermo Fisher Scientific). Band intensities were quantified using ImageJ software (NIH). NativeMark Unstained Protein Standard (Life Technologies) was used to estimate molecular weights of mitofusin protein complexes.

### Co-immunoprecipitation

Differentially tagged isolated mitochondrial populations (50 μg each) were mixed together. Mitochondria were incubated at 37°C for 30 minutes with beryllium fluoride (2.5 mM BeSO_4_, 25 mM NaF) with or without 2 mM GDP in fusion buffer (20 mM PIPES-KOH [pH 6.8], 150 mM KOAc, 5 mM Mg(OAc)_2_, 0.4 M sortibal with 0.12 mg/mL creatine kinase, 40 mM creatine phosphate, 1.5 mM ATP). Mitochondria were solubilized in lysis buffer (20 mM HEPES-KOH [pH 7.4], 50 mM KCl, 5 mM MgCl_2_) with 1.5% w/v n-Dodecyl β-D-maltoside (DDM), and 1X Halt Protease Inhibitor (Thermo Scientific) for 30 minutes on ice. Lysates were cleared at 10,000 x *g* for 15 minutes at 4°C. Supernatant was incubated with 50 mL magnetic µMACS Anti-DYKDDDDK MicroBeads (Miltenyi Biotec) for 30 minutes on ice. The sample was applied to a MACS Column (Miltenyi Biotec) placed in the magnetic field using a µMACS Separator (Miltenyi Biotec) and washed once with 300 mL 20 mM HEPES-KOH [pH 7.4], 50 mM KCl, 5 mM MgCl_2_, 0.1% DDM and once with 200 mL 20 mM HEPES-KOH [pH 7.4], 50 mM KCl, 5 mM MgCl_2_. One column volume (25 ml) SDS-PAGE loading buffer (60 mM Tris-HCl [pH 6.8], 2.5% sodium dodecyl sulfate, 5% βME, 5% sucrose, 0.1% bromophenol blue) was incubated for 15 minutes at room temperature and proteins were eluted once with 40 mL SDS-PAGE loading buffer. Samples were run on an SDS-PAGE gel and transferred onto nitrocellulose at 94V for 1 hour in 1X transfer buffer (25 mM Tris, 192 mM glycine, 20% methanol). Membranes were blocked in 4% Milk for at least 45 minutes and were probed with anti-Mfn1 antibody and anti-Mfn2 antibody for 4 hours at room temperature or overnight at 4°C. Membranes were incubated with DyLight secondary antibody (Invitrogen) at room temperature for 1 hour. Membranes were imaged on LI-COR Imaging System (LI-COR Biosciences).

### Western Blot Analysis

Protein lysates from MEFs were obtained by resuspending PBS washed cells in RIPA lysis buffer (150 mM NaCl, 1% Nonidet P-40, 1% Sodium deoxycholate, 0.1% SDS, 25 mM Tris [pH 7.4], 1X Halt Protease Inhibitor Cocktail, EDTA-Free [Thermo Scientific]). Samples were incubated on ice for 5 minutes and then spun at 21,000 x g for 15 minutes at 4°C. Supernatant was transferred to a clean tube and protein concentration was measured by BCA assay (Thermo Scientific). Samples were run on an SDS-PAGE gel and transferred onto nitrocellulose at 100 V for 50 minutes in 1X transfer buffer. Membranes were blocked in 4% milk for at least 45 minutes and were probed with anti-Mfn1, anti-Mfn2 (Sigma), anti-VDAC (Invitrogen) or anti-alpha Tubulin (Invitrogen) antibody for 4 hours at room temperature or overnight at 4°C. Membranes were incubated with DyLight secondary antibody (Invitrogen) at room temperature for 1 hour. Membranes were imaged on LI-COR Imaging System (LI-COR Biosciences).

### Bax expression and purification

Bax was purified as previously described (Suzuki et al. 2000). Briefly, pTYB1-Bax was expressed in E. coli strain BL21(DE3) grown in Luria-Bertani medium with 150 μg/mL ampicillin at 37°C to an OD600 ∼ 0.6, and protein expression was induced by the addition of 1 mM isopropyl 1-thio-β-D-galactopyranoside. Induced cultures were grown for about 3 hours at 37°C and then cells were harvested and frozen in liquid nitrogen and stored at −80°C. The pellet was resuspended in TEN buffer (20 mM Tris-HCl pH 8.0, 1 mM EDTA, 500 mM NaCl) and cells were lysed using a microfluidizer (Avestin). The lysate was subjected to centrifugation at 14,000 rpm for 45 min and the cleared lysate was passed through a 0.2 μm filter before binding to a chitin column equilibrated with TEN buffer. The column was washed with TEN buffer and then incubated with TEN buffer + 30 mM DTT for 48 hours at 4°C. Cleaved protein was eluted with TEN buffer and buffer exchange was performed with 20 mM Tris-HCl, pH 8.0 with 10% glycerol. Protein was bound to Q-sepharose and eluted with a linear gradient of NaCl. TCEP was added to Bax-containing fractions to a final concentration of 1 mM before the protein was aliquoted and frozen at −80°C.

### Plasmids & primers

The following plasmids were purchased from Addgene: pBABE-hygro (#1765), pBABE-puro (#1764), mito-PAGFP (#23348), pclbw-mito TagRFP (#58425), pclbw-mitoCFP (#58426). The following primers were used to for site directed mutagenesis by Gibson Assembly:

Mfn2^S378P^ F: (5’ – CCGTTCGTCTCATCATGGATCCCCTGCACATCGCAGC – 3’)

Mfn2^S378P^ R: (5’ – GCTGCGATGTGCAGGGGATCCATGATGAGACGAACGG – 3’)

Mfn2^A383V^ F: (5’ – ATTCCCTGCACATCGCAGTTCAGGAGCAGCGGG – 3’)

Mfn2^A383V^ R: (5’ – CCCGCTGCTCCTGAACTGCGATGTGCAGGGAAT – 3’)

Mfn2^Q386P^ F: (5’ –GCACATCGCAGCTCAGGAGCCGCGGGTTTATTGCCTAGAAATGCGG-3’)

Mfn2^Q386P^ R: (5’-CCGCATTTCTAGGCAATAAACCCGCGGCTCCTGAGCTGCGATGTGC-3’)

Mfn2^C390F^ F: (5’ – GGGTTTATTTCCTAGAAATGCGG – 3’)

Mfn2^C390F^ R: (5’ – CCGCATTTCTAGGAAATAAACCC – 3’)

Mfn2^L710P^ F: (5’ – GACATCACCCGAGATAATCCGGAGCAGGAAATTGCTGC – 3’)

Mfn2^L710P^ R: (5’ – GCAGCAATTTCCTGCTCCGGATTATCTCGGGTGATGTC – 3’)

## ACKNOWLEDGEMENTS

We thank the members of the Hoppins lab and Laura Lackner for critical scientific discussions and critical reading of the manuscript. We would also like to thank Richard Youle for sharing the pTYB1-Bax plasmid and Bax purification protocol. NBS was supported by National Institute of General Medical Sciences (NIH NIGMS) Training grant No. T32GM007270 and SCH is supported by NIH NIGMS Grant No. R01GM-118509.

## Notes

#### Summary of Updates

The updated manuscript includes new nucleotide dependent assembly data for Mfn2 and the hinge mutant variants. These data indicate that Bax enhances or stabilizes the 320 kDa oligomer observed in the presence of non-hydrolyzable analog of GTP. The hinge mutant variants respond to Bax in a manner similar to wild type, indicating that molecular defects in Mfn2 can be compensated for by cytosolic factors.

